# Diffusely abnormal white matter converts to T2 lesion volume in the absence of acute inflammation

**DOI:** 10.1101/2021.08.09.455717

**Authors:** M. Dadar, S. Mahmoud, S. Narayanan, DL. Collins, DL. Arnold, J. Maranzano

## Abstract

Diffusely abnormal white matter (DAWM), characterised by biochemical changes of myelin in the absence of frank demyelination, has been associated with clinical progression in secondary progressive MS (SPMS). However, little is known about changes of DAWM over time and their relation to focal white matter lesions (FWML).

The objectives of this work were: 1) To characterize the longitudinal evolution of FWML, DAWM, and DAWM that transforms into FWML, and 2) To determine whether gadolinium enhancement, known to be associated with the development of new FWML, is also related to DAWM voxels that transform into FWML.

Our data included 4220 MRI scans of 689 SPMS participants, followed for 156 weeks and 2677 scans of 686 RRMS participants, followed for 96 weeks. FWML and DAWM were segmented using a previously validated, automatic thresholding technique based on normalized T2 intensity values. Using longitudinally registered images, DAWM voxels at each visit that transformed into FWML on the last MRI scan as well as their overlap with gadolinium enhancing lesion masks were identified.

Our results showed that the average yearly rate of conversion of DAWM-to-FWML was 1.27cc for SPMS and 0.80cc for RRMS. FWML in SPMS participants significantly increased (t=3.9; p=0.0001) while DAWM significantly decreased (t=-4.3 p<0.0001) and the ratio FWML:DAWM increased (t=12.7; p<0.00001). RRMS participants also showed an increase in the FWML:DAWM Ratio (t=6.9; p<0.00001) but without a significant change of the individual volumes. Gadolinium enhancement was associated with 7.3% and 18.7% of focal New T2 lesion formation in the infrequent scans of the RRMS and SPMS cohorts, respectively. In comparison, only 0.1% and 0.0% of DAWM-to-FWML voxels overlapped with gadolinium enhancement.

We conclude that DAWM transforms into FWML over time, in both RRMS and SPMS. DAWM appears to represent a form of pre-lesional pathology that contributes to T2 lesion volume increase over time, independent of new focal inflammation and gadolinium enhancement.

## INTRODUCTION

Multiple Sclerosis (MS) is an inflammatory and neurodegenerative demyelinating disease characterized by different types of changes and lesions in the white and gray matter of the central nervous system (CNS)^1^. The majority of MS patients start with a relapsing-remitting course (RRMS) characterized by successive episodes of neurological dysfunction that partially or completely resolves ^1^. After a variable number of years of this relapsing course, the majority of patients experience a slow deterioration of neurological functions, unrelated to relapses, generally referred to as secondary progressive MS (SPMS) ^2^.

The MRI features of the earlier RRMS stage typically include the formation of new, focal white matter lesions (FWML) on T2-weighted scans. When they first develop, these new T2 lesions are generally associated with gadolinium (Gd) enhancement on T1-weighted images that lasts, on average, 3-4 weeks^3^. If the scanning frequency is adequate, most New T2 lesions are associated with Gd-enhancement^4 5^. If the interval between scans is 6-12 months, as was the case in our study, then only a portion of New T2 lesions will be associated with Gd-enhancement. New T2 lesions accumulate over time, becoming chronic. If chronic inactive, their volume stabilizes. If chronic active, their volume may show slow progressive expansion over time with progressively decreasing T1-weighted signal intensity associated with increasing tissue injury. All these different types of FWML appear as distinct areas of high signal intensity on T2w and fluid-attenuated inversion recovery (FLAIR) images, with sharp boundaries that clearly differentiate them from the surrounding tissue^4 5^.

Another type of WM hyperintensity visible on T2w images in MS involves regions that are less bright than FWML, but brighter than normal-appearing white matter (NAWM). Such regions generally have ill-defined borders and are referred to as diffusely abnormal white matter (DAWM)^6^. Histologic studies have shown that DAWM is characterized by a selective reduction of certain myelin phospholipids with relative preservation of myelin and axons. In particular, hemispheric slices of brain show a disproportionate reduction in myelin associated glycoprotein that corresponds to DAWM on MRI^7^. These observations point to a different pathophysiological substrate for DAWM compared to FWML. Additionally, quantitative MRI techniques (e.g. magnetization transfer ratio, relaxation times, and T1w-normalized intensities) have also established significant differences between areas of DAWM and FWML, with more severe alterations of MRI measures in FWMLs compared to DAWM^6^.

Previous studies have linked DAWM to disease progression as well as cortical pathology ^8, 9^. Our group has recently shown that SPMS participants exhibit DAWM transformation into FWML over time^10^ and that this transformation is significantly associated with clinical progression, making it a potential MRI marker of progressive biology in MS^10^. The few studies that have assessed changes of DAWM and FWML over time in RRMS populations have found that it rarely increases in this stage of MS^11^.

The present study takes advantage of a previously validated automated thresholding technique based on normalized T2 intensities^12^ to quantify the evolution of chronic FWML and DAWM volumes in RRMS and to compare them with the evolution in SPMS using two large longitudinal cohorts. We hypothesized FWML volume in MS (both RRMS and SPMS) accumulates *as a result of two distinct pathways*. The first, associated with a focal breach of the blood-brain-barrier (BBB) and acute extravasation of gadolinium, has been widely studied and used as a surrogate MRI marker of response to anti-inflammatory and immunomodulatory drugs^13^. The second pathway, which has not been directly demonstrated, would evolve by means of the slow transformation of DAWM into chronic FWML independent of acute focal inflammation. Hence, these transforming voxels would not exhibit Gd-enhancement, nor would they be expected to respond well to anti-inflammatory or immunomodulatory therapies. This second pathway would be, at least in part, responsible for the progression experienced by MS patients.

## METHODS

### Study Population

We included 4220 MRI scans of 689 SPMS participants, followed for three years (156 weeks) and scanned at screening and weeks 24, 48, 72, 96, 108, and 156, as well as 2677 MRI scans of 686 RRMS participants, followed for almost two years and scanned at screening, week 24, 48, and 96. Both cohorts were scanned with similar MRI acquisition protocols. All the participants had 3D T1w and 2D T2w images (Table 1) and were all followed-up consistently in the same scanner.

**Table 1:**
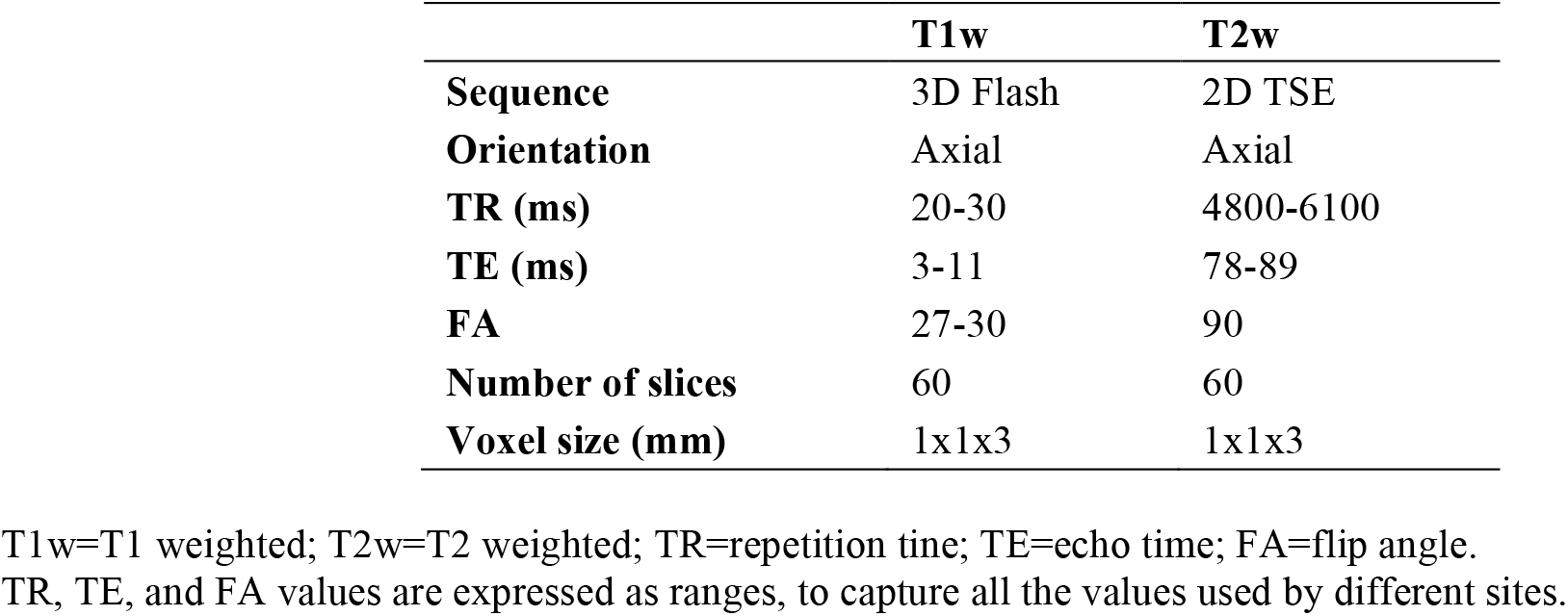
MRI acquisition parameters.

|  | **T1w** | **T2w** |
| --- | --- | --- |
| **Sequence** | 3D Flash | 2D TSE |
| **Orientation** | Axial | Axial |
| **TR (ms)** | 20-30 | 4800-6100 |
| **TE (ms)** | 3-11 | 78-89 |
| **FA** | 27-30 | 90 |
| **Number of slices** | 60 | 60 |
| **Voxel size (mm)** | 1x1x3 | 1x1x3 |

T1w=T1 weighted; T2w=T2 weighted; TR=repitition tine; TE=echo time; FA=flip angle. TR, TE, and FA values are expressed as ranges, to capture all the values used by different sites.

### Image Processing

All scans were processed using our standard image processing pipeline for longitudinal data, extensively described in our previous work^10, 12^. Briefly, the steps applied to the MRI data at each timepoint are: 1) Intensity non-uniformity correction^14^; 2) linear intensity normalization; 3) brain tissue extraction^15^; 4) tissue classification using T1w images^16^; 5) rigid co-registration of T2w to T1w images; 6) linear and nonlinear registration of the T1w images to MNI-ICBM152-2009c template^17, 18^; 7) automatic segmentation of WM lesions using a Bayesian Classifier, used to mask lesion voxels and obtain the intensity mode of the NAWM voxels^19^; 8) automatic separation of FWML from DAWM using intensity thresholds based on normalized T2 intensity mode values (the details of this automated separation techniques have been described in our previous studies)^10, 12^.

All the tools used in the image processing pipeline have been previously validated and used in multi-scanner datasets^10, 16, 17^. All the registrations and segmentation masks generated by the pipelines were visually evaluated and those that did not pass this quality control step (due to presence of artifacts such as motion, incidental findings, and registration failure) were excluded (n= 137 scans: 2%).

Once all timepoints were processed, a DAWM-to-FWML conversion mask was generated for each subject by identifying the DAWM voxels at baseline that changed to FWML at the last visit, using the longitudinally registered FWML and DAWM masks^10^. Finally, expert generated gadolinium-enhancing (Gd+) lesions and New T2 lesions masks ^20, 21^ from each timepoint were registered to the DAWM-to-FWML mask to quantify the percentage of overlap of Gd+ lesions with New T2 lesions, as well as the percentage of overlap of Gd+ lesions with DAWM-to-FWML voxels. We then excluded the Gd+ lesion and New T2 lesion voxels from the DAWM, FWML, and DAWM-to-FWML masks when quantifying the longitudinal changes of DAWM voxels to FWML in order to minimize any confounding effect of the large fluctuations in volume of acute inflammatory lesions. Our goal was to capture the longitudinal evolution of the chronic FWML and DAWM load that experiences slower and smaller volumetric changes, over time. Figure 1 shows two cases where FWML and DAWM have been identified.

**Figure 1:**
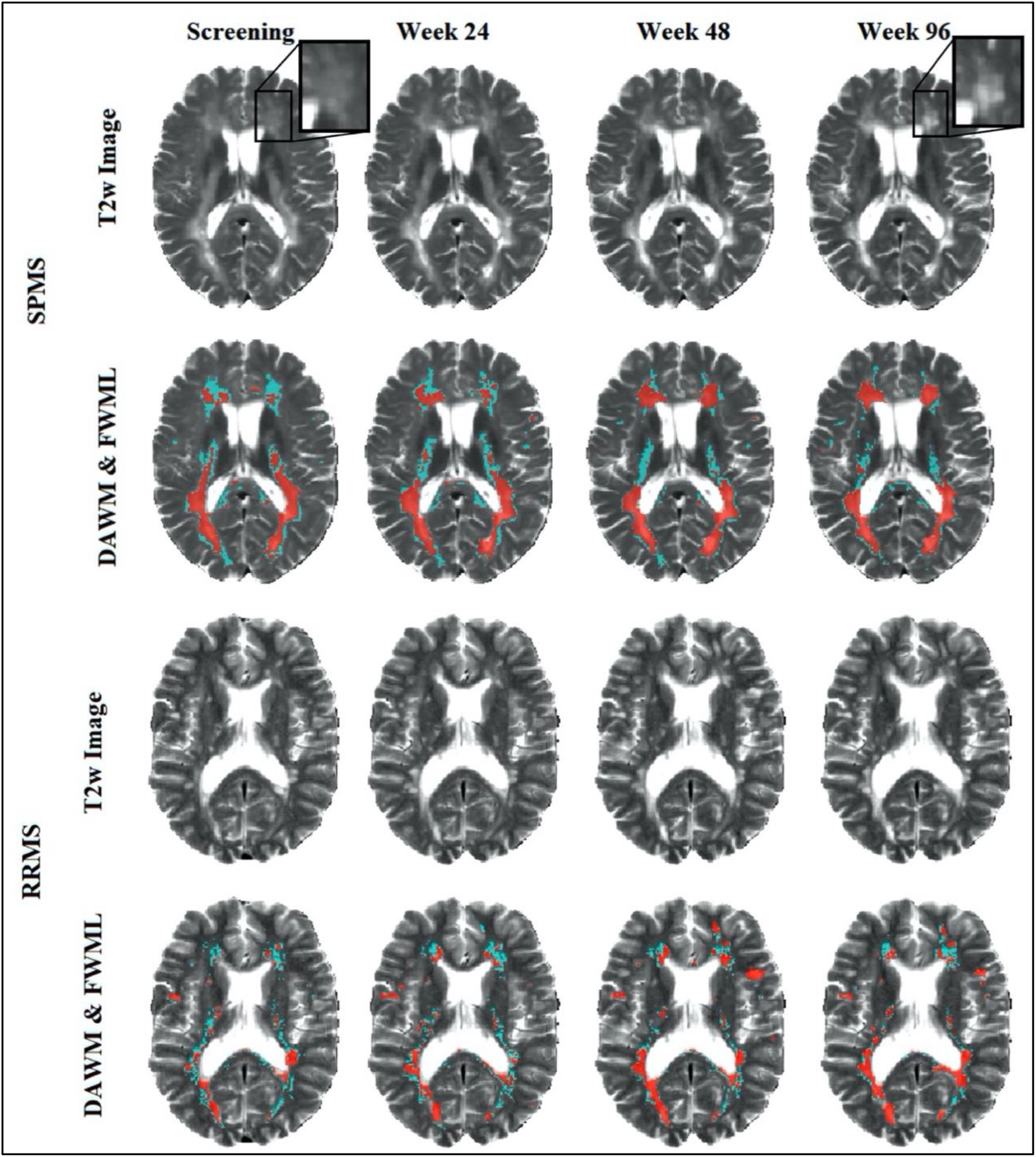
Example of two cases, one SPMS case (first two rows) and one RRMS case (bottom two rows) marking DAWM in green and FWML in red. Note the black boxes (zoomed-in first and last scans) where the transformation of DAWM into FWML is taking place. DAWM: Diffusely Abnormal White Matter. FWML: Focal White Matter Lesion.

### MRI variables of interest

We investigated changes in the following MRI variables of interest:

1. FWML volumes at each visit (excluding Gd+ and New T2)
2. DAWM volumes at each visit (excluding Gd+ and New T2)
3. FWML:DAWM-Ratio at each visit (excluding Gd+ and New T2)
4. Volume of DAWM at each visit that converted to FWML in the last visit (DAWM-to-FWML voxels, excluding Gd+ and New T2)
5. Rate of conversion of DAWM-to-FWML: volume of voxels that were DAWM at screening visit and FWML at last visit, divided by the duration between the two visits (i.e. 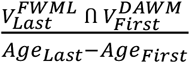)
6. Proportion of Gd+ lesions that overlap with NewT2 lesions at each visit (i.e. 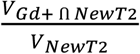)
7. Proportion of Gd+ lesions at each visit that overlap with DAWM-to-FWML voxels (i.e. 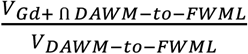)

### Statistical Analysis

We used descriptive statistics to present the demographic and MRI variables, as either mean and standard deviation (SD), or median and range, according to their distribution. Differences in sex proportion across RRMS and SPMS participants were explored using Chi-square, and differences in age at disease onset, age at first scan, disease duration, Expanded Disability Status Scale (EDSS)^22^, and FWML and DAWM volumes were assessed using Mann-Whitney U tests. Differences across disease types in volumes of DAWM and FWML at screening were also assessed using a generalized linear model, correcting for disease duration and age at disease onset. The longitudinal changes of DAWM and FWML volumes and FWML:DAWM ratios were assessed using linear mixed effects models, including age at disease onset, sex, and disease duration as independent variables, and considering subject and treatment as categorical random effects. A mixed effects model was also used to investigate whether the conversion rates of DAWM-to-FWML were significantly different between RRMS and SPMS groups, including age at onset, disease duration, and sex as covariates, and treatment as a random effect. All analyses were performed in SPSS version 26 and MATLAB version R2021a.

### Data availability

Derived data supporting the findings of this study are available from the corresponding author on request.

## RESULTS

The proportion of men and women and the median age at disease onset was not significantly different in RRMS and SPMS participants. As expected, the duration of the disease was significantly longer in SPMS participants. Table 2 presents all the demographic variables and the significances of the statistical tests used to explore the differences across groups.

**Table 2.** Demographic characteristics of the participants included in this study.

|  | **RRMS** | **SPMS** | **Significance** |
| --- | --- | --- | --- |
| **Number of participants** | 686 | 689 | n-a |
| **Sex** | 456 women (66.5%)<br>230 men (33.5%) | 425 women (61.7%)<br>264 men (38.3%) | 0.06 |
| **Age at the time of first scan (years)** | Median 38.0<br>Range 20.0 - 55.0 | Median 47.1<br>Range 23.9 - 58.0 | **<0.0001** |
| **Age at MS onset (years)** | Median 31.7<br>Range 18.0 - 53.5 | Median 30.6<br>Range 18.0 - 54.4 | 0.89 |
| **Disease duration (years)** | Median 4.5<br>Range 0.1 - 34.8 | Median 15.1<br>Range 2.0 - 39.9 | **<0.0001** |
| **EDSS** | Median 2.5<br>Range 0 - 6 | Median 6<br>Range 3 - 6.5 | **<0.0001** |

n-a=non-applicable; EDSS= Expanded Disability Status Scale

Median FWML volume at the first visit was significantly higher in SPMS (median: 13.76cc; IQR: 22.56) than RRMS participants (median 5.90cc; IQR: 12.55) using Mann-Whitney U test (p<0.0001) (Figure 3), and after correcting for age at disease onset and disease duration (p<0.0001). Median DAWM volume at the first visit was also significantly higher in SPMS (median: 23.47cc; IQR: 11.55) than RRMS participants (median 19.71cc; IQR: 11.94) using Mann-Whitney U test (p<0.0001), and after correcting for age at disease onset and disease duration (p=0.001). Figure 2 presents dual histograms showing the distribution of DAWM and FWML volumes for different levels of FWML and DAWM for both RRMS and SPMS.

**Figure 2:**
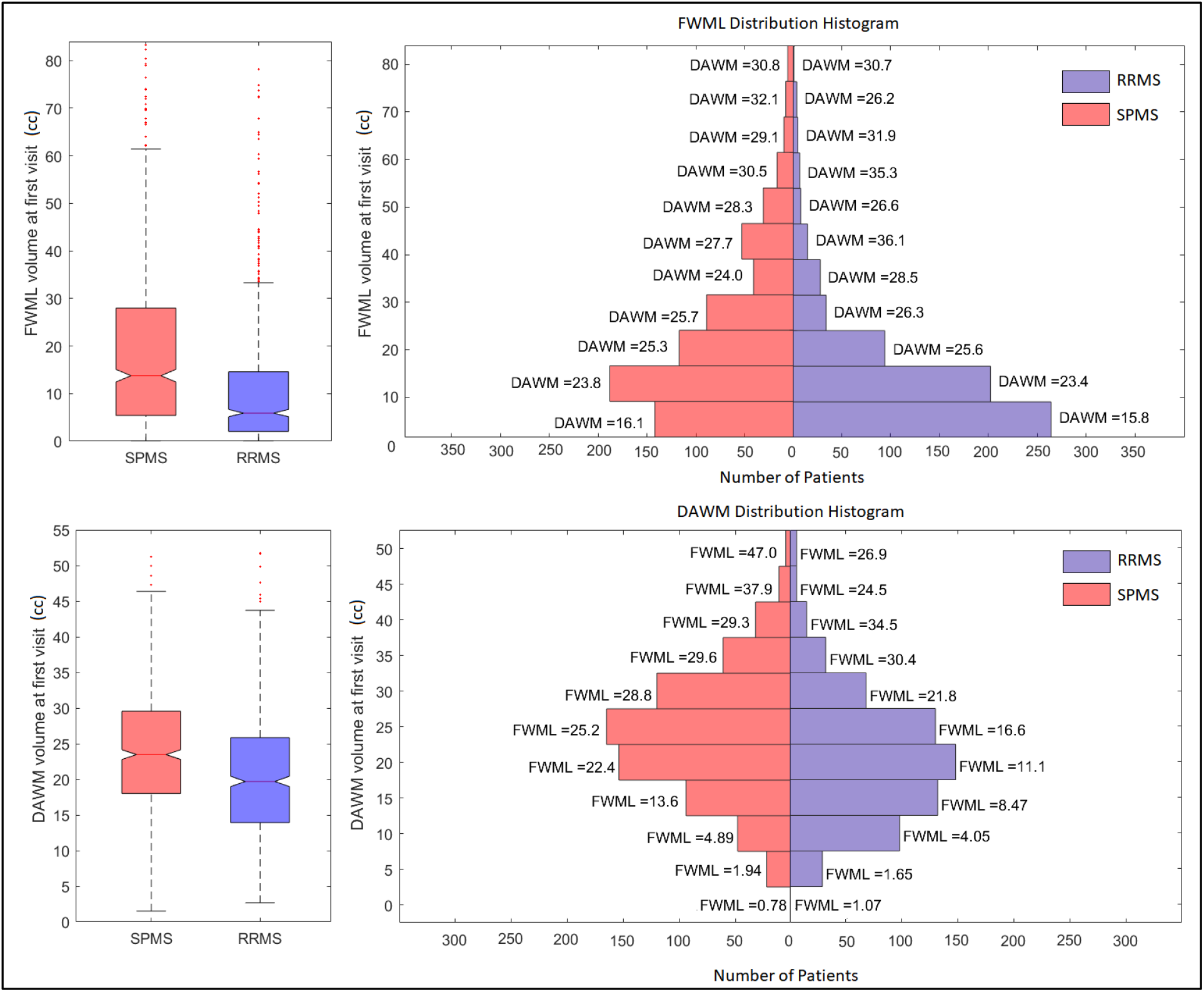
FWML and DAWM Volumes at first visit across groups (expressed in cubic centimeters). On the left: boxplots of FWML and DAWM volumes at first visit, separately for each group (SPMS in red, and RRMS in blue). On the right: histograms of FWML and DAWM volumes, along with mean volumes of DAWM for each FWML histogram bin, and vice-versa. FWML: Focal White Matter Lesion. DAWM: Diffusely Abnormal White Matter.

**Figure 3:**
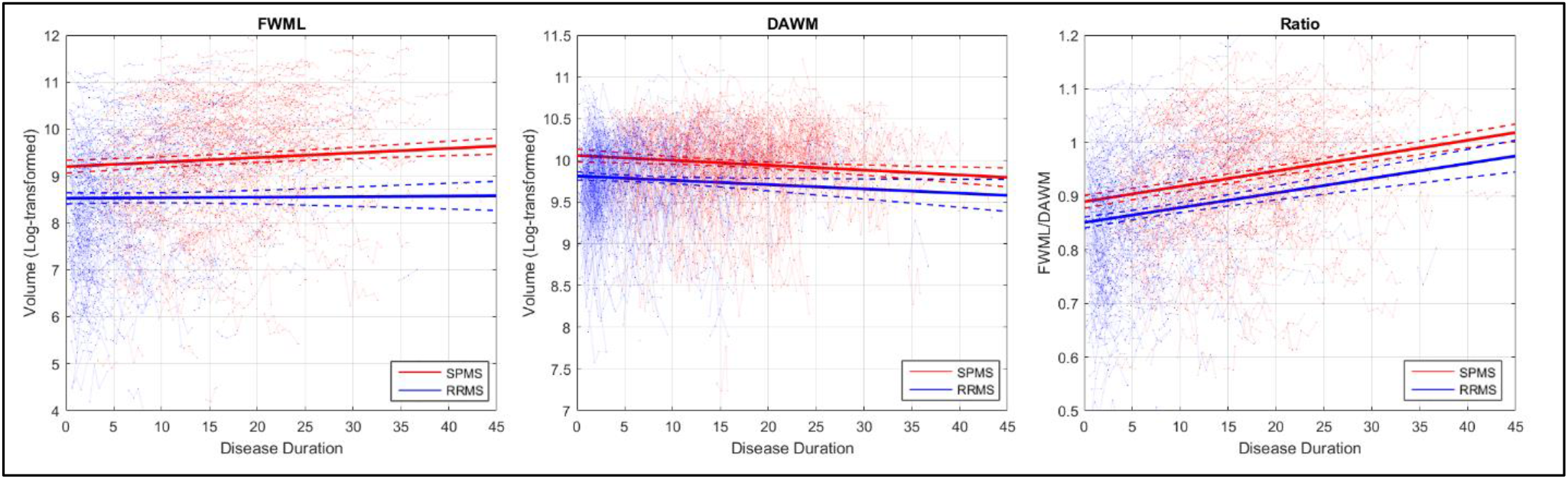
Regression graphs showing the volumes (log-transformed) of FWML, DAWM and the FWML/DAWM Ratio in relation to the duration of the disease. FWML: Focal White Matter Lesion. DAWM: Diffusely Abnormal White Matter.

Figure 3 shows the longitudinal changes in FWML volumes, DAWM volumes, and their ratios in RRMS and SPMS populations. SPMS participants showed volumes of FWML that significantly increased (t=3.9; p=0.01) along with a significant decrease of DAWM volumes (t=-4.3; p<0.0001), as well as a significant FWML:DAWM Ratio increase (t=12.7; p<0.00001). RRMS participants showed a significant increase in the FWML:DAWM Ratio (t=6.9; p<0.00001), however changes in FWML and DAWM volumes were not individually significant in RRMS. Note that treatment was not associated with DAWM, FWML, or FWML:DAWM volumes in either group.

The yearly average DAWM-to-FWML conversion rates were 1.27cc and 0.80cc for SPMS and RRMS, respectively. The mixed effects model also showed that this difference was statistically significant (t = 3.98, p <0.0001). Finally, Gd+ voxels showed 7.3% and 18.7% overlap with new focal lesions (New T2 lesions) in RRMS and SPMS, respectively. In comparison, only 0.1% and 0.0% of DAWM-to-FWML voxels overlapped with Gd+ lesions in RRMS and SPMS, respectively.

## DISCUSSION

Using longitudinally registered MRIs, we have shown that voxels of DAWM progressively convert to chronic FWML (i.e. focal lesions not associated with Gd+ or New T2 lesions) over time in both RRMS and SPMS patients. The yearly rate of this transformation was significantly higher in SPMS than in RRMS, but was present in RRMS as well, consistent with other evidence that disability progression ^23^ and progressive biology ^24^ is not exclusive to progressive MS. This has important implications for the understanding of how T2w lesion volume accumulates in MS, indicating that it does not necessarily accumulate only as a result of the residual abnormality from acute, focal inflammation. To support this hypothesis, we assessed the co-localization of Gd+ lesions with two different types of T2 hyperintense lesions: new focal lesions, also known (and named throughout this manuscript) as New T2 lesions, which are a marker of acute inflammatory demyelination, and the conversion of DAWM to FWML. New T2 lesions showed 7.3% and 18.7% overlap with Gd+ enhancing lesions in RRMS and SPMS groups, respectively, whereas voxels that converted from DAWM-to-FWML showed negligible (0.1 and 0.0%) enhancement.

The nature of DAWM on MRI is not well understood. Histopathology of DAWM shows biochemical abnormalities of myelin associated glycoprotein (MAG) that extend beyond the regions of obvious abnormality of other histochemical markers indicative of demyelination^7^. This is consistent with earlier histochemical observations of Itoyama et al.,^25^ reporting decreases in MAG immunostaining extending far beyond the margins of acute demyelination and affecting myelin sheaths that appeared otherwise normal. The authors suggested that the abnormality of MAG preceded myelin breakdown, and that, since MAG is synthesized in the oligodendroglia and localized to the periaxonal membrane, the decreased MAG might be associated with an eventual failure to maintain the integrity of the myelin sheath. They and others ^26, 27^ have suggested that loss of MAG may be a marker of metabolic stress, in particular oxidative stress. More recently, Metz et al.^28^ have correlated preferential loss of MAG with pattern III lesions, which have an ill-defined lesion edge, are less likely to show contrast enhancement than are other MS lesion types, and are associated with a distal oligodendrogliopathy, also related to oxidative stress. Other stresses, such as abbreviated exposure to cuprizone, may also lead to subtle biochemical changes to myelin in the absence of overt demyelination, and to subsequent inflammatory demyelination^29^. Taken together, these observations support the pathophysiological concept that an early myelinopathy present in DAWM may be the prelude to overt demyelination.

The last decades have been marked by significant advances in the treatment and monitoring of RRMS using MRI to quantify changes in the formation of new lesions as a marker of treatment response^13, 30^. Gd+ lesions and development of New T2 lesions have been particularly useful for predicting the response of clinical relapses to new anti-inflammatory therapies. However, these types of lesions reflect the response of the acute, peripherally mediated inflammatory component of the disease. Unfortunately, changes in the brain that may relate more directly to the biology associated with disability progression in the absence of relapses ^23^ are more difficult to identify and quantify. The conversion of DAWM to FWML may be important in this regard. It has been previously suggested that T2 lesions may form in the absence of gadolinium enhancement^31^, but the mechanisms that might be responsible for this phenomenon were not evident. Our analysis supports the concept that a slowly progressive increase in T2 lesion volume may be driven by the conversion of DAWM to chronic FWML, and that this phenomenon is not exclusive to SPMS, but is also present in RRMS. We have previously shown that DAWM-to-FWML transformation is more pronounced in SPMS patients that have confirmed disability progression for 12 and 24 weeks^11^. We now extend the findings of this transformation to RRMS, where disability progression independent of relapses is increasingly appreciated to also occur^23^. The volume of tissue undergoing DAWM-to-FWML transformation suggests that, along with other ongoing pathology^32^, this transformation may play a significant role determining progression. Further studies quantifying different aspects of progressive biology including DAWM conversion to FWML, slowly expanding lesions^24^, phase (or paramagnetic) rim lesions^33^, cortical demyelination^34^ and the effects of inflammation on normal-appearing white matter are necessary to better understand the relative contribution to progression of these different pathologies.

Our study is not without limitations. Our automatic classifier of DAWM and FWML was developed based on expert segmentations of MRI in the absence of a histological gold standard. However, the image processing pipeline generates robust and consistent results, and it has been previously validated in test-retest settings and used in multi-scanner datasets^11^. Our RRMS cohort is a distinct group, different than the SPMS group, and included patients in very different stages of the disease. Ideally, the same group of RRMS patients should be prospectively followed through the transitional phase up to the secondary progressive phase to unequivocally determine that the change from RRMS to SPMS goes in parallel with a change in the amount of DAWM-to-FWML transformation. To partially address this issue, we selected two groups that were followed using similar and consistent MRI acquisition protocols. Additionally, the subjects included were on a variety of treatments that were effective in suppressing acute new lesion formation. This means that the acute component in these cohorts (i.e. the Gd+ lesions and New T2 lesions) was much less than what would be expected during the natural course of the un-treated disease. Another limitation regards the presence of atrophy. To ensure that CSF signal due to atrophy was not included in any of the masks, we used tissue segmentations at each timepoint^16^ and detected hyperintensities (both DAWM and FWML) within white matter masks that were eroded by one voxel, avoiding the periventricular adjacent voxels where lesion voxels could be contaminated by CSF signal related to ventricular enlargement. This approach, intended to avoid the confounding effect of atrophy, provided a conservative quantification of the longitudinal increase in FWML and transformation of DAWM-to-FWML.

## CONCLUSION

DAWM is a prelesional state that transforms into FWML over time in MS. This transformation occurs in the absence of gadolinium-enhancement and likely contributes significantly to increases in T2 lesion volumes that occur in progressive MS in the absence of ongoing focal inflammatory lesion activity.

## FUNDING

This Investigation was supported in part by an award from the International Progressive MS Alliance, award reference number PA-1603-08175. MD is supported a scholarship from the Canadian Consortium on Neurodegeneration in Aging as well as an Alzheimer Society Research Program (ASRP) postdoctoral award.

## AUTHOR CONTRIBUTIONS

**Mahsa Dadar:** Study concept and design, analysis and interpretation of the data and drafting and revising the manuscript.

**Sawsan Mahmoud**: Analysis of the data and revising the manuscript.

**Sridar Narayanan:** Interpretation of the data and revising the manuscript.

**Douglas L. Arnold:** Interpretation of the data and revising the manuscript.

**D. Louis Collins:** Interpretation of the data and revising the manuscript.

**Josefina Maranzano:** Study concept and design, analysis and interpretation of the data and drafting and revising the manuscript.

## CONFLICTS OF INTEREST

Dr. Dadar has nothing to disclose.

Sawsan Mahmoud has nothing to disclose.

Dr. Narayanan reports grants from the Canadian Institutes of Health Research, the Myelin Repair Foundation and the International Progressive MS Alliance, during the conduct of the study; personal fees from NeuroRx Research, personal fees from Genentech, and travel assistance from MedDay, outside the submitted work.

Dr. Arnold reports grants from Canadian Network of MS Clinics, during the conduct of the study; personal fees from Acorda, personal fees from albert Charitable Trust, personal fees from Biogen, personal fees from Celgene, personal fees from Roche, personal fees from GeNeuro, personal fees from Frequency Therapeutics, personal fees from MedDay, personal fees from Merck Serono, personal fees from Novartis, personal fees from Sanofi-Aventis, personal fees from Wave Life Sciences, grants from Biogen, grants from Immunotec, grants from Novartis, other from NeuroRx, outside the submitted work.

Dr. Collins reports grants from Canadian Institute of Health Research, during the conduct of the study; personal fees from NeuroRx inc, outside the submitted work.

Dr. Maranzano has nothing to disclose.

## Abbreviations

BBB: Blood Brain Barrier
CNS: Central Nervous System
CSF: CerebroSpinal Fluid
DAWM: Diffusely Abnormal White Matter
FLAIR: FLuid-Attenuated Inversion Recovery
FWML: Focal White Matter Lesions
Gd: Gadolinium
Gd+: Gadolinium enhancing
IQR: Inter-Quartile Range
MRI: Magnetic Resonance Imaging
MS: Multiple Sclerosis
NAWM: Normal Appearing White Matter
RRMS: Relapsing Remitting Multiple Sclerosis
SPMS: Secondary Progressive Multiple Sclerosis
WM: White Matter
MAG: Myelin Associated Glycoprotein

